# A monogenean parasite reveals the widespread translocation of the African Clawed Frog in its native range

**DOI:** 10.1101/2021.10.21.465306

**Authors:** Anneke L Schoeman, Louis H du Preez, Nikol Kmentová, Maarten PM Vanhove

## Abstract

1. The management of bio-invasions relies upon the development of methods to trace their origin and expansion. Co-introduced parasites, especially monogenean flatworms, are ideal tags for the movement of their hosts due to their short generations, direct life cycles and host specificity. However, they are yet to be applied to trace the intraspecific movement of host lineages in their native ranges.
2. As proof of this concept, we conducted a comparative phylogeographic analysis based upon two mitochondrial markers of a globally invasive frog *Xenopus laevis* and its monogenean parasite *Protopolystoma xenopodis* in its native range in southern Africa and invasive range in Europe.
3. Translocation of lineages was largely masked in the frog’s phylogeography. However, incongruent links between host and parasite phylogeography indicated host switches from one host lineage to another after these were brought into contact in the native range. Thus, past translocation of host lineages is revealed by the invasion success of its co-introduced parasite lineage.
4. This study demonstrates that parasite data can serve as an independent line of evidence in invasion biology, also on the intraspecific level, shedding light on previously undetected invasion dynamics. Based upon the distribution of these invasive parasite lineages, we infer that there is widespread anthropogenic translocation of this frog, not only via official export routes, but also facilitated by the frog’s use as live bait by angling communities.
5. *Synthesis and applications.* Data from co-introduced, host-specific parasites, as tags for translocation, can add value to investigations in invasion biology and conservation. A better understanding of the translocation history and resulting genetic mixing of host and parasite lineages in the native range can shed light on the genetic make-up of parasite assemblages co-introduced to the invasive range. Knowledge of the intraspecific movement of different lineages of animals in their native ranges also has conservation implications, since contact between divergent lineages of hosts and parasites can facilitate host switches and altered parasite dynamics in both native and invasive populations. Therefore, we recommend the inclusion of parasite data as a more holistic approach to the invasion ecology of animals on the intraspecific level.

## 1 INTRODUCTION

The Anthropocene is characterised by the unparalleled human-mediated passage of organisms from their natural ranges to ecosystems across the world (Capinha *et al*., 2015; Ricciardi, 2007). If these non-native organisms manage to disperse from their initial introduction point and seed new populations in their non-native range, they are regarded as invasive (Blackburn *et al*., 2011; Richardson *et al*., 2011). This reshuffling of the world’s biota is a decisive agent of global change, as evidenced by the fact that invasive species are implicated as the cause of a third of recent animal extinctions (Blackburn *et al*., 2019).

Clearly, the development of methods to trace both the origin and expansion of biological invaders is an urgent task. Genomic tools can be informative for the reconstruction of invasion histories, the prediction of potential range expansion and invasion success, and the development of control strategies (e.g. de Busschere *et al*., 2016; Dufresnes *et al*., 2017; Dlugosch & Parker, 2008; Hiller & Lessios, 2017; Hudson *et al*., 2022; Lee & Gelembiuk, 2008; Prentis *et al*., 2009; Stouthamer *et al*., 2017; Zahiri *et al*., 2019). Nonetheless, in many cases the available molecular data or chosen molecular markers are of insufficient resolution by themselves to trace the movement of species (e.g. Ahyong *et al*., 2017; Criscione *et al*., 2006). Thus, despite the accessibilty of genomic tools and the widespread use of phylogeographic analyses in invasion science, these methods certainly have their limitations (Fitzpatrick *et al*., 2012; Rius & Turon, 2020).

A more holistic approach to solving the origin and range shifts of invasive species involves molecular data from co-introduced parasites (Gagne *et al*., 2021). Such an integrative approach has been proposed to investigate invasions that are difficult to trace (Clavero *et al*., 2016). For example, the long-debated cryptogenic history of the European marine snail *Littorina littorea* (Linnaeus, 1758) was solved with the help of an associated digenean (Platyhelminthes: Digenea) parasite (Blakeslee *et al*., 2008). Likewise, intraspecific morphometric variation of monogenean flatworm (Platyhelminthes: Monogenea) parasites identified both the origin and introduced life stage of invasive clupeid fish in Africa (Kmentová *et al*., 2019), shed light on the history of an invasive population of odontobutid fish in Europe (Ondračková *et al*., 2012) and provided information on the introduction route of invasive gobiid fish in Europe (Huyse *et al*., 2015). Moreover, the genetic variation among monogenean gill parasites were used to illuminate the introduction history of non-native cichlid fish in Africa (Geraerts *et al*., 2022; Jorissen *et al*., 2022).

Matters are complicated even further when we wish to trace movement of conspecific organisms between different populations. This phenomenon, also known as intraspecific cryptic invasion, describes the process where one lineage of a species spreads into parts of the species’ native range where another lineage is already present, often with human help (Morais & Reichard, 2018; Saltonstall, 2002). Once again, co-translocated parasites hold great promise to provide information on the movement of lineages of their hosts. Monogenean parasites in particular exhibit sufficient levels of phylogeographic structuring to assess intraspecific spatial patterns in their hosts (Baldwin *et al*., 2012; Hahn *et al*., 2015; Huyse *et al*., 2017; Kmentová *et al*., 2018). At the very least, the phylogeographic structuring of monogenean lineages can support evidence revealed by the host data set or historic records, such as the well-documented anthropogenic introduction of salmonid fish host stocks across Europe that is reflected in the phylogeography of their gyrodactylid monogenean lineages (Hahn *et al*., 2015). Monogenean parasites are regarded as ideal tags of host genealogy due to their short generations, direct life cycles and generally high host-specificity (Nieberding & Olivieri, 2007; Pariselle *et al*., 2011). Yet, the comparative phylogeographic analysis of host and parasite data has never been applied to trace intraspecific cryptic invasions within the native ranges of species.

The African Clawed Frog *Xenopus laevis* (Daudin, 1802) (Anura: Pipidae) is arguably the world’s most widely distributed and well-studied amphibian (Measey *et al*., 2012; van Sittert & Measey, 2016). Previous studies on the morphology and genetics of *X. laevis* have identified several differentiated populations in South Africa that are segregated by large mountain ranges and different climatic regions, with the most notable divergence between northern and southern populations (de Busschere *et al*., 2016; du Preez *et al*., 2009; Furman *et al*., 2015; Grohovaz *et al*., 1996; Measey & Channing, 2003). In opposition to the clear phylogeographic structuring stands the widespread occurrence of both natural and human-mediated translocation of *X. laevis* in the region. This movement is a consequence of naturally occurring overland migration, the domestic export of the frog for teaching and learning and the widely adopted practice of the translocation of the frog for use as live bait in recreational angling (de Villiers & Measey, 2017; Measey & Davies, 2011; Measey, 2016; Measey *et al*., 2017; van Sittert & Measey, 2016; Weldon *et al*., 2007). One would expect these translocations to leave a genetic trace (van Sittert & Measey, 2016). Yet, the cryptic invasion of *X. laevis* over long distances and across major geographic barriers in the native range has not been confirmed in previous phylogeographic studies (de Busschere *et al*., 2016; du Preez *et al*., 2009; Furman *et al*., 2015; Measey & Channing, 2003).

We posit that the bladder polystomatid flatworm *Protopolystoma xenopodis* (Price, 1943) Bychowsky, 1957 (Monogenea: Polystomatidae) is the ideal tool to trace the hitherto unconfirmed contact between northern and southern lineages of its host *X. laevis* across its native range of southern Africa. In natural populations across sub-Saharan Africa, *P. xenopodis*, the most widely distributed of the species of *Protopolystoma*, is restricted to a single host species group, the *leavis* group, even in sympatry with other *Xenopus* species (Tinsley & Jackson, 1998a,b). Furthermore, in *P. xenopodis*, notable intraspecific morphological variation exists between populations of the polystomatid infecting *X. laevis* in southern Africa and those infecting more northerly host species (Tinsley & Jackson, 1998b). This pattern is echoed even on a smaller regional scale in South Africa, where notable morphological and phylogenetic divergence exists between northern and southern populations of *P. xenopodis* that parasitise *X. laevis* (Schoeman *et al*., 2022). The combination of high host specificity and intraspecific divergence suggests that *P. xenopodis* could be useful in a holobiont approach to a bio-invasion study, since any inferred movement of the parasite has to be mediated by movement of the host.

Therefore, in the present study, we use the widely distributed and intimately associated *X. laevis* and *P. xenopodis* system as a proof of concept that intraspecific parasite phylogeography can provide information on the cryptic introductions of their host lineages. We hypothesise that the phylogeography of *P. xenopodis* in South Africa will magnify that of its host *X. laevis*, which will allow us to employ a comparative phylogeographic analysis of this host and parasite in the context of their domestic translocation. We postulate that the parasite phylogeography will reveal contact between distinct lineages of *X. laevis* where evidence for translocation events might be lacking in the frog phylogeography and provide new insights on invasion pathways in the native range and beyond.

## 2 MATERIALS AND METHODS

### 2.1 Host and parasite sampling

In the native range of southern Africa, 46 adult *X. laevis* were captured from March 2017 to April 2019 in liver baited funnel traps from 30 localities (Fig. 1; see Appendix S1 for locality details). The localities were representative of the phylogeography of the frog, based upon the lineage assignment of de Busschere *et al*. (2016) (see Appendix S2 for locality classification). In the invasive range, 13 adult *X. laevis* were collected from 12 localities in western France and one in Portugal during the summers of 2017, 2018 and 2019 (see Appendix S1 for locality details). In France, frogs were captured in baited traps by members of the EU LIFE CROAA team as part of the regional eradication programme for invasive amphibians. Frogs in Portugal were obtained from the locally captured laboratory stock of the University of Lisbon.

**FIGURE 1.**
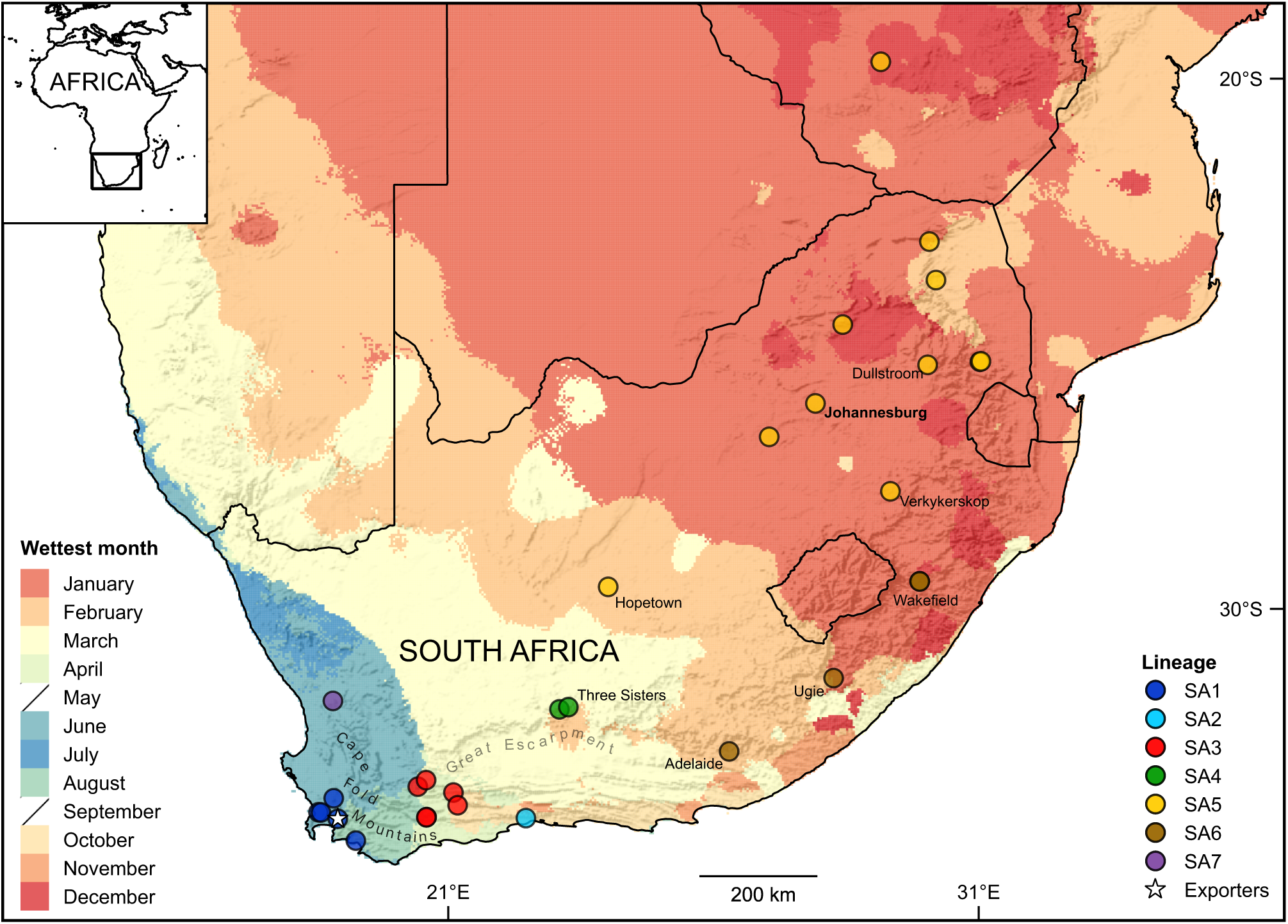
Localities in southern Africa where 46 adult *Xenopus laevis* were collected. Localities are classified according to the geographic extent of the native phylogeographic lineages of *X. laevis* (de Busschere *et al*., 2016). Key localities are named according to the closest town. Jonkershoek (A) is the source locality of the majority of historic *X. laevis* exports. Mountain ranges are indicated with shaded relief and rainfall seasonality is represented by the wettest month of each 25 km^2^ grid cell. Monthly mean precipitation data, obtained from the WorldClim 2.1 database at a spatial resolution of 2.5 arc-minutes (Fick & Hijmans, 2017), were used to identify the months with the highest historical mean precipitation from 1970 to 2000 for each grid cell. Elevation data were obtained from the Natural Earth database. The map was constructed in QGIS *version* 3.10.2-A Coruña (QGIS Development Team, 2018) with the Mercator projection.

All frogs underwent double euthanasia with anaesthesia in 6% ethyl-3-aminobenzoate methanesulfonate (MS222) (Sigma-Aldrich Co., USA) followed by pithing as required by the institutional ethics committee. During subsequent dissection, 59 *P. xenopodis* (one per frog) were retrieved from the bladders for molecular analyses. Polystomatids and liver tissue from their corresponding frog hosts were preserved in 70% ethanol.

### 2.2 Molecular data collection

For the comparative phylogeographic analyses, we obtained the genetic sequences of the mitochondrial barcoding genes for cytochrome *c* oxidase subunit I (*COX1*) and for the small 12S subunit of ribosomal RNA (*rrnS*) of both the frogs and the polystomatids. The DNA was extracted from host and parasite tissue with the PCRBIO Rapid Extract PCR kit (PCR Biosystems Ltd., UK) or the *Quick*-DNA^TM^ Miniprep Plus kit (Zymo Research, USA). Subsequently, amplification reactions for the mitochondrial *COX1* and *rrnS* genes of both frogs and polystomatids were prepared with 2 to 5 *µ*L genomic DNA, 1.25 *µ*L forward primer [0.2 *µ*M], 1.25 *µ*L reverse primer [0.2 *µ*M], 12.5 *µ*L master mix and PCR-grade water to the final volume of 25 *µ*L. The 2× PCRBIO HS Taq Mix Red (PCR Biosystems Ltd., UK) or the OneTaq^®^ 2× Master Mix with Standard Buffer (New England Biolabs Inc., USA) were used with the temperature of the elongation steps as per supplier instructions (68 °C or 72 °C).

For the frog *COX1*, the ‘LCO1490’/’HCO2198’ (Folmer *et al*., 1994) and ‘VF1’/’VR1’ (Ivanova *et al*., 2006; Ward *et al*., 2005) primer pairs were implemented with the following thermocycling profile: initial denaturation at 95 °C for 2 min, followed by 35 cycles of denaturation at 95 °C for 30 s, annealing for 30 s at 50 °C for the first primer pair and 54 °C for the second and elongation for 2 min, terminated by a final elongation step of 7 min. In the case of the ‘12SA-F’/’12SB-R’ primer pair (Kocher *et al*., 1989) to obtain the frog *rrnS*, the profile of Amaro *et al*. (2013) was followed. For the amplification of the polystomatid *COX1* and *rrnS*, the profile of Héritier *et al*. (2015) was implemented for the primer pairs ‘L-CO1p’/’H-Cox1R’, ‘L-CO1p’/’H-Cox1p2’, ‘12SpolF1’/’12SpolR1’ and ‘12SpolF1’/’12SpolR9’ (Héritier *et al*., 2015; Littlewood *et al*., 1997).

For purification and sequencing, PCR products were sent to a commercial company (Inqaba Biotec, RSA) that used the ExoSAP protocol (New England Biolabs Ltd.) for purification and obtained the sequences with BigDye^®^ Terminator *version* 3.1 Cycle Sequencing, utilising the corresponding primer pairs used in the PCR reaction, on an ABI3500XL analyser. Sequences were assembled and manually edited in Geneious Prime *version* 2021.0.3 (www.geneious.com). The 236 obtained sequences were deposited in GenBank (accession numbers to be added after manuscript revision).

### 2.3 Calculation of sequence statistics

An investigation of sequence diversity is an exploratory analysis that can reveal the relative differentiation within and between geographic clusters. It can also give an indication of the markers’ applicability to subsequent analyses. For these analyses, we obtained alignments of each of the four gene sets: *COX1* or *rrnS* of the frog and *COX1* or *rrnS* of the polystomatid. The Clustal Omega algorithm under default settings (Sievers & Higgins, 2017) implemented in Geneious Prime was used for the alignment of all four data sets. For the protein-coding *COX1* gene, translation alignment was implemented under the vertebrate mitochondrial genetic code (table 2) for the frog and the echinoderm and flatworm mitochondrial genetic code (table 9) for the polystomatid.

Genetic diversity indices were calculated for all specimens combined and for geographic clusters in the host and parasite samples. Number of haplotypes and polymorphic sites, haplotype diversity, nucleotide diversity and mean percentage of pairwise differences were calculated in Arlequin *version* 3.5 (Excoffier & Lischer, 2010). Three groups were defined based upon collection locality (see Appendix S2 for full details), namely specimens collected in South Africa to the southwest of the Great Escarpment, the major geographical barrier to dispersal in the region (SA1, SA2, SA3, SA4 and SA7; n = 30), to the northeast of the Great Escarpment (SA5 and SA6; n = 16), or in their non-native range in France (n = 12). The specimen from Portugal was not included as a separate group in the analyses because of small sample size.

### 2.4 Phylogenetic analyses

The phylogenetic relationships between the different populations of both the host *X. laevis* and the parasite *P. xenopodis*, represented in phylogenetic trees derived from the *COX1* and *rrnS* genes, formed the basis of the subsequent comparative phylogeographic analysis.

The sequences were prepared for the phylogenetic analyses through removal of duplicates, addition of outgroups for rooting, re-alignment and concatenation. Duplicate sequences were identified with the help of CD-HIT (Fu *et al*., 2012; Li & Godzik, 2006) implemented online (Huang *et al*., 2010). Only specimens with unique *COX1* or *rrnS* gene haplotypes were retained for the analysis, leading to 30 unique sequence combinations (23 unique haplotypes of *COX1* and 10 of *rrnS*) in the case of *X. laevis* and 44 (34 of *COX1* and 35 of *rrnS*) in the case of *P. xenopodis*. In order to root the trees, the *COX1* and *rrnS* gene sequences of the congeneric Mü ller’s clawed frog *X. muelleri* (Peters, 1844) (GenBank accession numbers EU599031, AY581657; Evans *et al*., 2004, Evans *et al*., 2008) and its polystomatid *P. occidentalis* Tinsley *et* Jackson, 1998 (GenBank accession numbers KR856179, KR856121; Héritier *et al*., 2015) were added as outgroups. This polystomatid only infects species from the *muelleri* group, even in sympatry with *X. laevis* (Tinsley & Jackson, 1998b). Furthermore, its host *X. muelleri* reportedly reproduces sterile offspring when hybridisation occurs between this species and its sympatric congener *X. laevis* (Fischer *et al*., 2000). Therefore, the selected outgroups are applicable to an intraspecific analysis of the *X. laevis* and *P. xenopodis* host-parasite system. The resulting data sets, after removal of duplicates and addition of the outgroups, were re-aligned by gene as detailed above. The two aligned genes of each species were concatenated by individual in Geneious Prime. Ultimately, 31 sequences of a total of 1153 bp (717 bp *COX1* and 436 bp *rrnS*), containing 189 sites with gaps and 183 variable sites, of which 88 were parsimony informative, were used for the phylogenetic analysis of the frog data set. Likewise, the 45 aligned sequences consisting of a total of 934 bp (423 bp *COX1* and 511 bp *rrnS*), with 799 complete sites, of which 225 polymorphic sites with a subset of 174 parsimony informative sites, were use to infer the phylogeny of the parasite.

Prior to phylogenetic inference, the most optimal models of molecular evolution of each gene partition were determined. Initially, the concatenated alignments comprised four partitions each, namely the three codon positions of the partial *COX1* gene and the *rrnS* gene. The ModelFinder selection routine (Kalyaanamoorthy *et al*., 2017), in conjunction with the FreeRate heterogeneity model (Soubrier *et al*., 2012), implemented in IQ-TREE *version* 2.1.2 (Minh *et al*., 2020), was employed to select the optimal model of molecular evolution for each partition. To reveal the optimal partitioning scheme and most suitable evolutionary models for each alignment, the initial partitions were sequentially merged (Chernomor *et al*., 2016) until model fit, as evaluated by the Bayesian Information Criterion (BIC), ceased to improve.

Finally, we obtained phylogenies of the host and the parasite using the maximum likelihood (ML) approach. The ML phylogeny of the host was inferred under the TIM2 model (Posada, 2003) with edge-unlinked FreeRate heterogeneity models (Soubrier *et al*., 2012), proportions of invariant sites (Gu *et al*., 1995) and discrete gamma models (Yang, 1994) for all partitions. Similarly, the ML tree of the polystomatid was inferred under the TPM3 model (Kimura, 1981) with a FreeRate heterogeneity model (Soubrier *et al*., 2012) for all partitions. The analyses were performed in IQ-TREE *version* 2.1.2 (Minh *et al*., 2020), with the assessment of branch support through ultrafast bootstrapping (UFboot2; Hoang *et al*., 2018) and the Shimodaira-Hasegawa approximate likelihood ratio test (SH-aLRT; Guindon *et al*., 2010), each with 10 000 replicates. The two phylogenies were rooted with respect to the outgroups in TreeGraph2 *version* 2.15.0-877 *beta* (Stöver & Müller, 2010). Clades with UFboot2 support lower than 95% or SH-aLRT lower than 80% were collapsed to polytomies in the same programme.

### 2.5 Comparative phylogeographic analyses

The assessment of the global fit between the two phylogenies at the level of lineages provided a framework to interpret the mismatches or incongruencies represented by certain individual host-parasite links as indications of long distance movement and subsequent parasite switching to different host lineages. The global congruence of the two phylogenies was assessed by the implementation of both Procrustean Approach to Co-phylogeny (PACo) (Balbuena *et al*., 2013) and the fourth-corner statistical approach of ParaFit (Legendre *et al*., 2002). Both these methods consider the contribution of individual specimen interactions to phylogenetic congruence (Balbuena *et al*., 2013; Legendre *et al*., 2002; Poisot & Stouffer, 2015). Moreover, both methods can accommodate trees that are not fully resolved and support instances of uneven numbers of terminal nodes of hosts and parasites.

In PACo, we assessed the dependency of the parasite phylogeny upon that of the host through asymmetrical Procrustean superimposition of host and parasite distance matrices, whilst ParaFit tested the dependence of both phylogenies upon one another (Balbuena *et al*., 2013; Legendre *et al*., 2002). Prior to analyses, the outgroups were removed from the ML phylogenies. The phylogenies were converted to matrices of patristic distances by the package *ape* (Paradis & Schliep, 2019) in R *version* 4.0.2 (R Core Team, 2020). These matrices served as input for both analyses, along with the corresponding host-parasite association matrix, where the polystomatids were assigned to their original host individuals, or to a host with an identical gene sequence to the original host individual.

The principal coordinates for PACo were derived with the R package *paco* (Hutchinson *et al*., 2017) *version* 0.4.2 for the Procrustes superimposition in conjunction with the Cailliez correction (Cailliez, 1983) to avoid the production of negative eigenvalues of non-Euclidean phylogenetic distances. The PACo analysis was performed, using the asymmetric Procrustes statistic to test for phylogenetic tracking by the parasite of its host, in *paco* with 10 000 permutations, with conserved row sums and number of interactions. The dynamics of the interacting clades were further explored by identifying those interactions that most contribute to the possible concordance between the two phylogenies through a jackknife procedure in *paco*. Those interactions with the smallest residual distances were interpreted as stronger supporters of overall congruence, whilst those with larger residual distances represented likely mismatches between host and parasite phylogeny.

The ParaFit analysis (Legendre *et al*., 2002) was carried out with *ape* based upon the patristic distance matrices, using 10 000 permutations with the Cailliez correction (Cailliez, 1983). Both the ParaFitLink tests were employed to assess the significance of the individual links (Legendre *et al*., 2002), which also pointed out host-parasite links that signify incongruencies.

Visualisation was achieved through a tanglegram of the host and parasite ML phylogenies with a weighted interaction network, based upon the residuals yielded by the PACo jackknife procedure. It was constructed with the R package *phytools* (Revell, 2012) with the function *cophylo()*, which has an internal algorithm that rotates nodes to optimally match tips. Due to the removal of duplicate sequences, certain host and parasite terminal nodes were involved in multiple interactions.

Haplotype genealogies based upon bifurcarting phylogenetic trees and Fitch (1970) distances between sequences are an alternative way of visualising phylogenies whilst giving an indication of the number of individual specimens represented by each unique sequence in the data set. For this reason, we also visualised the 59 *X. laevis* and the 59 *P. xenopodis*, that is with the inclusion of duplicates, in two haplotype genealogies. The sequences were aligned and concatenated as detailed above. Two ML phylogenies, one for the frog and one for the parasites, based on all 59 sequences of each and no outgroups were derived using the procedure described above. These trees, along with the re-aligned sequences, served as input for Fitchi (Matschiner, 2015), which produced two haplotype genealogy graphs.

### 2.6 Use of frogs by anglers

As an addition to the information provided by (Weldon *et al*., 2007) on the use of *X. laevis* as live bait by anglers in South Africa, we obtained anecdotal information on fishing practices from several stakeholders whilst conducting our fieldwork across the region. We asked about the practices surrounding the use of *X. laevis* as live bait during fishing and the translocation and release of *X. laevis* during fishing activities. Nine recreational and subsistence anglers in the Eastern Cape, Limpopo, Northwest, KwaZulu-Natal and Gauteng Provinces provided information on the use of *X. laevis* as live bait by their communities in the vicinity of some of our sampling localities.

## 3 RESULTS

### 3.1 Sequence diversity

The genetic diversity of the two parasite genes was generally higher than the two orthologous host genes, with greater genetic diversity among the samples collected in South Africa than in France in both host and parasite (Table 1). Genetic diversity was generally higher in the hosts collected to the southwest of the Great Escarpment than in those collected from the northeast, which was not the case for the parasites (Table 1).

**TABLE 1.** Genetic diversity of the bladder flatworm *Protopolystoma xenopodis* and its frog host *Xenopus laevis*, grouped according to sampling locality – collected in South Africa to the southwest or to the northeast of the Great Escarpment, or in France. Indices were inferred separately by gene for each group and for all samples combined from the aligned partial mitochondrial *COX1* and *rrnS* genes.

| | *n* | H | L | S | Hd | $\pi$ | d (%) |
| --- | --- | --- | --- | --- | --- | --- | --- |
| *X. laevis COX1* |  |  |  |  |  |  |  |
| Southwestern SA | 30 | 11 | 641 | 78 | 0.823 ± 0.061 | 0.026 ± 0.013 | 2.6 |
| Northeastern SA | 16 | 6 | 641 | 74 | 0.742 ± 0.084 | 0.025 ± 0.013 | 2.5 |
| France | 12 | 3 | 641 | 50 | 0.591 ± 0.108 | 0.014 ± 0.008 | 1.4 |
| All specimens | 58 | 16 | 641 | 86 | 0.880 ± 0.023 | 0.048 ± 0.024 | 4.8 |
| *X. laevis rrnS* |  |  |  |  |  |  |  |
| Southwestern SA | 30 | 7 | 369 | 13 | 0.662 ± 0.088 | 0.008 ± 0.005 | 0.8 |
| Northeastern SA | 16 | 3 | 369 | 2 | 0.342 ± 0.140 | 0.001 ± 0.001 | 0.1 |
| France | 12 | 2 | 369 | 5 | 0.303 ± 0.148 | 0.004 ± 0.003 | 0.4 |
| All specimens | 58 | 9 | 369 | 14 | 0.681 ± 0.044 | 0.009 ± 0.005 | 0.9 |
| *P. xenopodis COX1* |  |  |  |  |  |  |  |
| Southwestern SA | 30 | 20 | 416 | 94 | 0.959 ± 0.022 | 0.070 ± 0.035 | 7.0 |
| Northeastern SA | 16 | 15 | 416 | 89 | 0.992 ± 0.025 | 0.072 ± 0.037 | 7.2 |
| France | 12 | 3 | 416 | 3 | 0.591 ± 0.108 | 0.004 ± 0.003 | 0.4 |
| All specimens | 58 | 34 | 416 | 110 | 0.958 ± 0.014 | 0.072 ± 0.035 | 7.2 |
| *P. xenopodis rrnS* |  |  |  |  |  |  |  |
| Southwestern SA | 30 | 14 | 484 | 78 | 0.903 ± 0.036 | 0.050 ± 0.025 | 5.0 |
| Northeastern SA | 16 | 14 | 484 | 87 | 0.983 ± 0.028 | 0.055 ± 0.029 | 5.5 |
| France | 12 | 2 | 484 | 1 | 0.485 ± 0.106 | 0.001 ± 0.001 | 0.1 |
| All specimens | 58 | 28 | 484 | 98 | 0.930 ± 0.020 | 0.048 ± 0.024 | 4.8 |

**Abbreviations:** SA, South Africa; *n*, sample size; H, number of haplotypes; L, length of sequence; S, number of polymorphic sites; Hd, haplotype diversity; $\pi$ , nucleotide diversity; d, mean percentage of pairwise differences.

### 3.2 Phylogenetic relationships of host and parasite

The phylogenetic analyses confirmed marked phylogeographic structuring in both the host and parasite in their native range, which provided the necessary resolution to assess spatial patterns. Tree reconstruction showed that frogs from the northeastern SA5 and SA6 localities in the native range form a well-supported clade, along with some of the frogs from the SA4 locality (clade NE in Fig. 2 with UFboot2 = 95% / SH-aLRT = 95.8%). In addition, two clades with unresolved relationships to one another and to NE could be identified among sampled frogs from the southwestern localities in the native range of *X. laevis* (clades SW_coast_ with UFboot2 = 81% / SH-aLRT = 86.1% and SW_inland_ with UFboot2 = 98% / SH-aLRT = 99.7% in Fig. 2). On the whole, members of the first clade originate from localities closer to the coastline (clade SW_coast_ in Fig. 2 from SA1, SA2, SA3, SA4 and SA7 localities) than the second one (clade SW_inland_ in Fig. 2 from SA3, SA4 and SA6 localities). The non-native frog from the invasive population in Portugal, along with one of the non-native frogs from western France, grouped together with the native frogs in SW_coast_ in the host phylogeny, thus placing these invasive frogs in the same clade as the frog from Jonkershoek (Fig. 1), the source of the majority of *X. laevis* exports. Nonetheless, the invasive population of *X. laevis* in France also harboured several frogs that clustered with frogs from the native northeastern localities in the NE clade.

**FIGURE 2.**
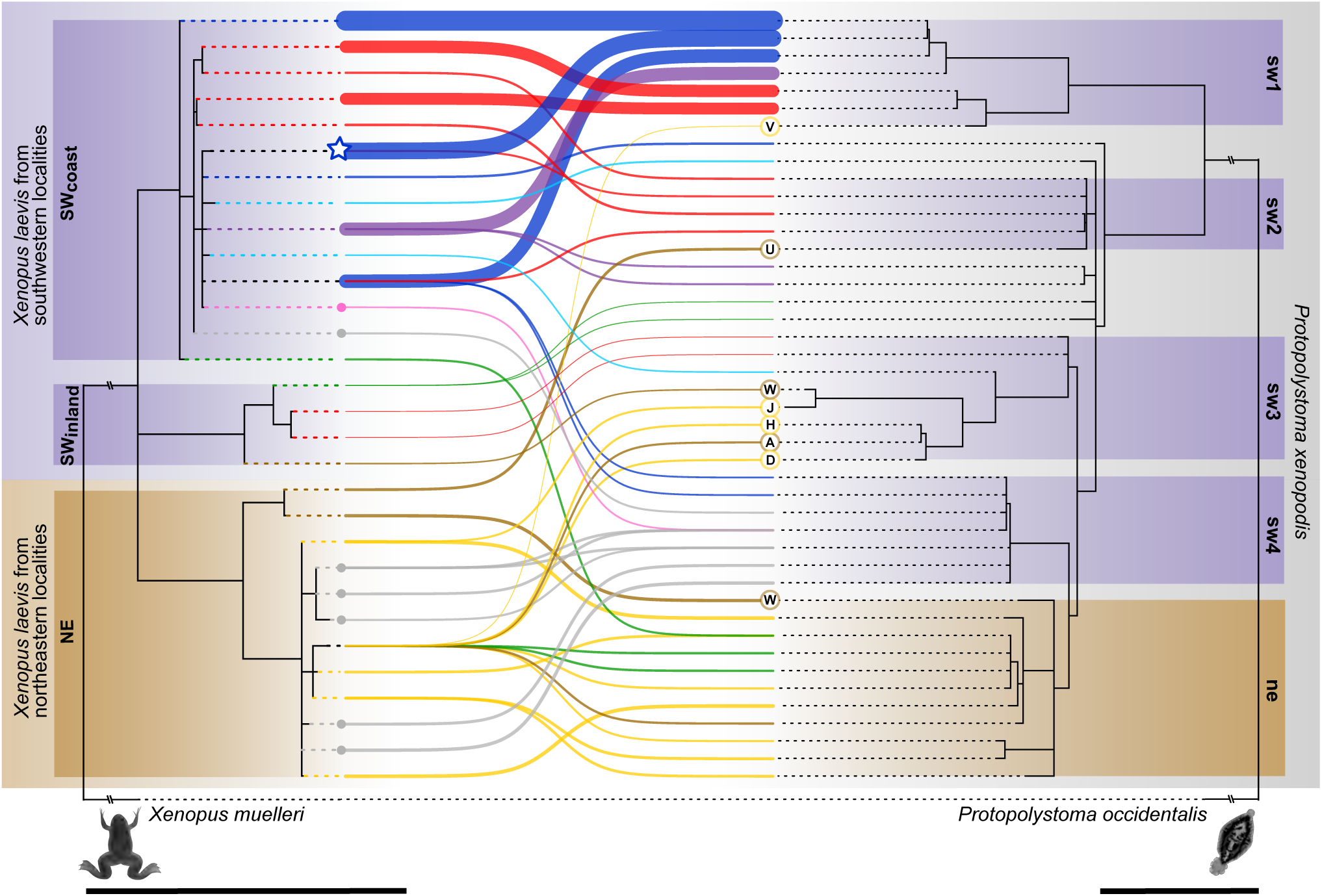
The interaction network and phylogenies of the frog *Xenopus laevis* (left) and its polystomatid *Protopolystoma xenopodis* (right) with *X. muelleri* and *P. occidentalis* as outgroups. Topologies are based on maximum likelihood analyses of concatenated *COX1* and *rrnS* gene alignments of 31 frogs and 45 polystomatids with low support clades (UFboot2 *<* 95% or SH-alrt *<* 80%) collapsed. Links indicate observed host-parasite associations, weighted according to the contribution of each interaction to the overall phylogenetic congruence with thicker lines corresponding to greater support. Colours refer to geographic origin to the southwest (SW) and northeast (NE) of the Great Escarpment, following the classification of de Busschere *et al*. (2016): SA1 (dark blue), SA2 (light blue), SA3 (red), SA4 (green), SA5 (yellow), SA6 (brown) and SA7 (purple) (see Fig. 1) with main lineages named and highlighted. Some parasite specimens from southwestern SA1, SA2, SA3, SA4 and SA7 localities are not assigned to any of the main lineages due to their paraphyletic grouping basal to lineages **sw3**, **sw4** and **ne**. Invasive *X. laevis* and co-introduced *P. xenopodis* from France (grey) and Portugal (pink) are also included. Key localities are Adelaide (A), Dullstroom (D), Hopetown (H), Johannesburg (J), Ugie (U), Verkykerskop (V) and Wakefield (W). Jonkershoek (A) is the origin of the majority of *X. laevis* exports. Scale bars indicate 0.1 nucleotide substitutions per site.

In the same manner as the host phylogeny, the ML inference of the *P. xenopodis* tree revealed a single, well-supported clade of polystomatids from hosts from the northeastern SA5 and SA6 localities, along with some of the parasites from the SA4 locality (clade **ne** in Fig. 2 with UFboot2 = 97% / SH-aLRT = 87.6%). Several clades basal to the **ne** group emerged among the polystomatids from hosts collected at the southwestern SA1, SA2, SA3, SA4 and SA7 localities (clades **sw1**, **sw2**, **sw3** and **sw4** in Fig. 2). The earliest diverging clade is a well-supported grouping of polystomatids collected from SA1, SA3 and SA7, along with one polystomatid from SA5 (clade **sw1** in Fig. 2 with UFboot2 = 94%/SH-alrt = 88.1%). Two other native southwestern clades included a clade of polystomatids from the SA3 localities and one from SA6 (clade **sw2** in Fig. 2 with UFboot2 = 53%/SH-alrt = 90.7%) and another clade of polystomatids from southwesten SA2 and SA3 localities and some northeastern SA5 and SA6 localities (clade **sw3** in Fig. 2 with UFboot2 = 61% / SH-aLRT = 86.6%). Another well-supported clade, the sister clade to **ne**, contained polystomatids from SA1, France and Portugal (clade **sw4** in Fig. 2 with UFboot2 = 100% / SH-aLRT = 100%). However, overall low support values of many internal nodes in the parasite ML tree, especially the nodes basal to the **sw3**, **sw4** and **ne** lineages, hampered the interpretation of phylogenetic clustering among some of the polystomatids collected from the southwestern localities. Consequently, some parasite specimens were not assigned to any clades.

### 3.3 Congruence of host and parasite phylogeographies

The phylogeny of *X. laevis* exhibited significant overall congruence with that of *P. xenopodis* (PACo 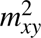 = 0.1034, *p* = 0.0018, n = 10000; ParaFitGlobal = 0.0202, *p <* 0.0001, n = 10000), with the PACo analysis suggesting that the phylogeny of *P. xenopodis* tracked that of *X. laevis*. According to PACo, links between frogs from SW_coast_ and their corresponding polystomatids that fall within **sw1** in the parasite phylogeny, generally contributed the most to the overall phylogenetic congruence. The establishment of overall congruence between these two phylogenies allowed us to hone in on mismatches between the two phylogenies, which signified past contact between divergent clades.

Based upon visual inspection of the tanglegram (Fig. 2), a few links represented mismatches between the host and parasite phylogenies that could be explained by long distance translocation of frogs from the SW host lineages to the northeastern localities followed by spill-over of **sw1**, **sw2** and **sw3** polystomatids to NE hosts. These indications of host switching were identified in frog-polystomatid pairs from SA5 and SA6 localities (Dullstroom, Hopetown, Johannesburg, Verkykerskop, Adelaide, Ugie and Wakefield), where NE hosts harboured parasites that clustered with their southwestern **sw1**, **sw2** and **sw4** counterparts. The PACo jackknife procedure further identified the interactions between frog–polystomatid pairs from the invasive range in France and Portugal as links conveying low support to congruence. Moreover, interactions between frogs and polystomatids from the contact zone near Three Sisters (SA4) and Wakefield (SA6) demonstrate low support to overall congruence. Finally, the ParaFitLink tests reported 30 out of 48 links as significant at a level of 0.05. Out of the 18 links that did not significantly support congruence, 11 signified host switches by parasites to another host lineage that occurred in the invasive range of France and Portugal and at SA4, SA5 and SA6 localities in the native range (Johannesburg, Three Sisters, Ugie and Wakefield).

The haplotype genealogies of the frog host and the polystomatid parasite (Fig. 3) unsurprisingly demonstrated the same phylogeographic structuring as those revealed by the phylogenetic analysis of *X. laevis* (Fig. 2). The frog haplotype genealogy clearly revealed that all northeastern specimens clustered together, except for frog specimens from Wakefield (SA5), which clustered with both the NE and SW_inland_ clades, and the frogs collected at Three Sisters (SA4), which were present in all three host lineages. This situation was somewhat different for the polystomatids, where the haplotype genealogy illustrated that the cluster of parasite specimens from northeastern hosts (**ne**) contained only nine of the sixteen specimens collected from the northeastern SA5 and SA6 localities. The seven remaining parasite specimens from SA5 and SA6 localities (Adelaide, Dullstroom, Hopetown, Johannesburg, Ugie, Verkykerskop and Wakefield) were spread across the remainder of the network, therefore clustering with polystomatids from southwestern hosts (**sw1**, **sw2**, **sw3**). Moreover, parasites collected at the Three Sisters (SA4) localities clustered with both northeastern (**ne**) and southwestern (**sw2**) parasites.

**FIGURE 3.**
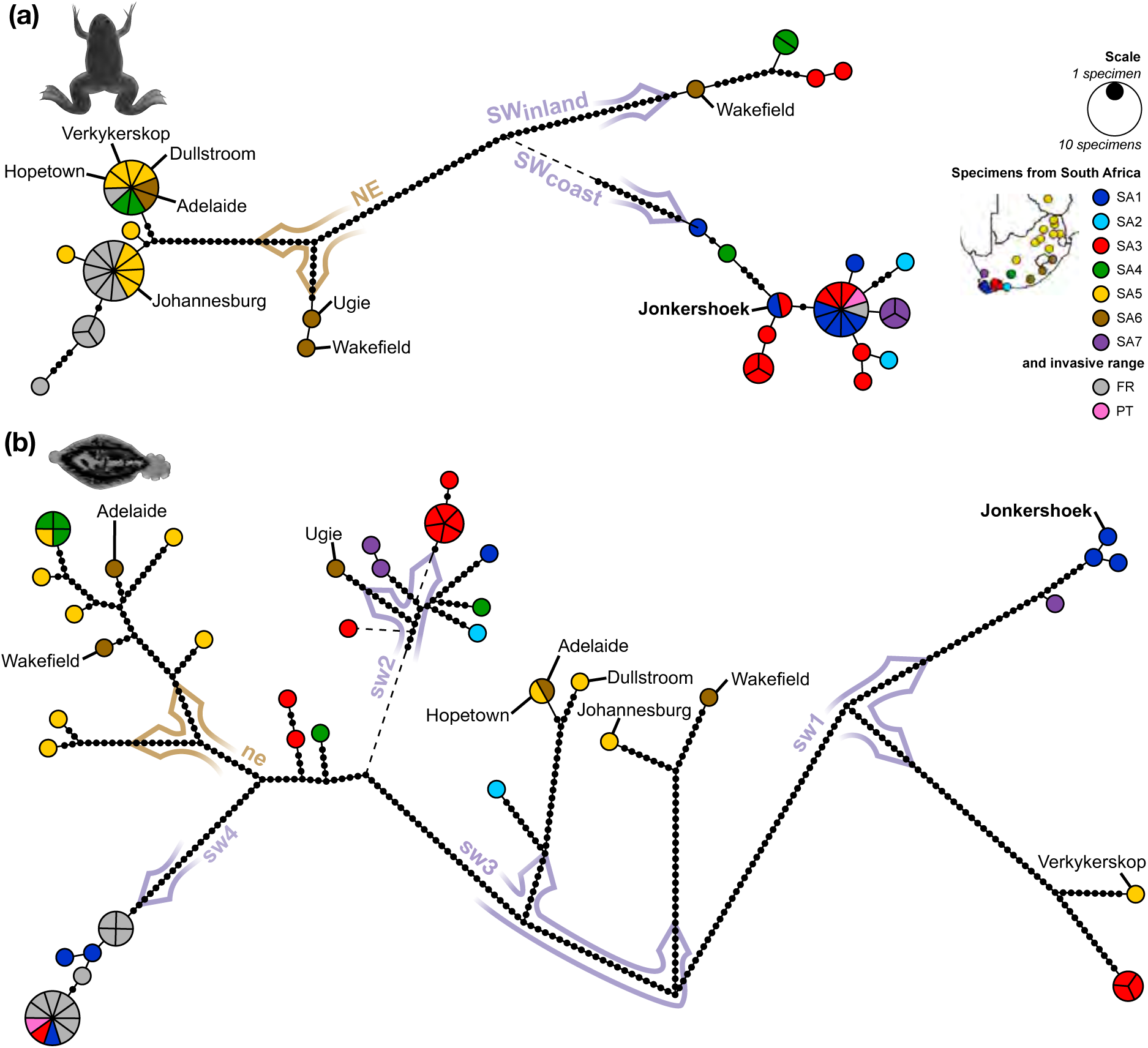
Unrooted haplotype genealogy graphs of the partial *COX1* and *rrnS* genes of **(a)** 59 *Xenopus laevis* specimens and **(b)** their corresponding 59 *Protopolystoma xenopodis* parasites. For frogs and their polystomatids collected in the native southern Africa (see inset map), colours refer to the phylogeographic lineages of the host (de Busschere *et al*., 2016). Non-native *X. laevis* and *P. xenopodis* are from France (grey) and Portugal (pink). Highlighted haplotype groups correspond to the clades revealed by the phylogenetic analyses (Fig. 2): northeastern host (NE) and parasite (ne) and southwestern host (SW) and parasite (sw) clades. Key localities with host lineage switches are named, as well as Jonkershoek, the source locality of most of the *X. laevis* exports. Sizes of circles are proportional to haplotype frequencies (see scale). The number of mutational steps is marked on the connecting branches with dots. Dashed lines are extensions of dotted sections for improved visualisation.

### 3.4 Translocation of frogs by anglers

Our informal discussions with anglers confirmed the widespread use of *X. laevis* as live bait in five out of the nine South African provinces, with all nine anglers confirming the existence of the practice in their communities and surrounding areas. The frog was usually employed in the capture of black bass *Micropterus* Lacepède, 1802 and African sharp-toothed catfish *Clarias gariepinus* (Burchell, 1822). Black bass are non-native inhabitants of water bodies across South Africa (Hargrove *et al*., 2015; Khosa *et al*., 2019) and African sharp-toothed catfish are non-native to all areas to the south of the Great Escarpment (Weyl *et al*., 2016). Seven out of the nine anglers indicated that specimens of *X. laevis* are generally caught and used locally, with short range translocation of the captured specimens. However, two of the anglers admitted to buying *X. laevis* from breeders and pet shops in urban areas (Potchefstroom and Johannesburg) and transporting them to more rural areas for use as live bait. These fishermen were active near some of our sampling localities, namely Potchefstroom, Johannesburg, Dullstroom and Verkykerskop. All fishermen confirmed that surplus live *X. laevis* are routinely released on site and that escapees are common. Notably, a fisherman in Ugie described the practice of acquiring live *X. laevis* from water bodies in the vicinity and translocating specimens to stock local dams for future use.

## 4 DISCUSSION

In this study, the widespread, yet previously hidden, domestic expansion of *X. laevis* lineages in South Africa is brought to light through a comparison with the phylogeography of its co-invasive parasite, *P. xenopodis*. The use of parasite data to reveal host connectivity is hardly novel (reviewed in Gagne *et al*., 2021). Yet, the present study is the first application of host-parasite phylogeographic analyses to the detection of intraspecific cryptic invasions. Furthermore, it demonstrates that even a comparatively modest data set of less than 60 host-parasite pairs can reveal compelling evidence for historic translocation.

### 4.1 Parasite clustering mirrors the intraspecific phylogeography of the host

If the phylogeographic structuring of a parasite mirrors that of its host, a phenomenon which is not necessarily a consequence of co-evolutionary processes (de Vienne *et al*., 2013), the genealogy of the parasite can provide insight on the translocation of the host (Nieberding & Olivieri, 2007). Based on our findings, there is phylogenetic congruence between the African Clawed Frog and its monogenean parasite. This means that the observed phylogeographic structuring in the parasite, which is supported by both molecular and morphological evidence (Schoeman *et al*., 2022), can corroborate and expound upon the phylogeography of the host *X. laevis*, which also shows notable phylogeographic structuring across its native range (de Busschere *et al*., 2016; du Preez *et al*., 2009; Furman *et al*., 2015; Grohovaz *et al*., 1996; Measey & Channing, 2003). After a brief free-swimming stage, *P. xenopodis* completes the rest of its life cycle in the excretory system of *X. laevis* (Theunissen *et al*., 2014), which translates to a very specific and intimate host-parasite association that can faithfully portray the phylogeographic structure of the frog. However, the parasite data set provides an independent line of evidence and not a mere echo of the phylogeographic structuring of the frog, since each parasite represents a separate infection event from a pool of free-swimming larvae.

The genetically distinct host-parasite clusters display distributions that roughly follow the two climatic regimes in the region, namely the summer rainfall region to the northeast and the winter rainfall region to the southwest, with the plateau edge of the Great Escarpment and the many ridges of the Cape Fold Mountains as assumed geographical barriers to natural dispersal (Furman *et al*., 2015). The more pronounced clustering among hosts and parasites and the higher haplotype diversity in hosts that hail from the southwest of the Great Escarpment is likely brought about by the presence of the many ridges of the Cape Fold Mountains (de Busschere *et al*., 2016; du Preez *et al*., 2009; Furman *et al*., 2015; Grohovaz *et al*., 1996; Measey & Channing, 2003). On the other hand, well-documented human-mediated translocation of *X. laevis* in the area to the southwest of the Great Escarpment (van Sittert & Measey, 2016; Weldon *et al*., 2007), in conjunction with widespread overland migration of the frog with the help of the many interconnected artificial water bodies in the region (de Villiers & Measey, 2017; Measey, 2016), ensure some gene flow – and exchange of parasites – between the divergent frog populations in this area.

### 4.2 Parasite spillover is an indication of contact between divergent host lineages

Parasite phylogeography offers a tool to recover historical insights of host dispersal pathways, since host lineage switches reflect past interactions between host organisms at various time scales (e.g. Barbosa *et al*., 2012; Criscione *et al*., 2006; Galbreath & Hoberg, 2015; Galbreath *et al*., 2020; Nieberding *et al*., 2004). The usefulness of parasite species in reflecting the introduction pathways of their hosts has been demonstrated for fish hosts and their monogenean parasites on a few occasions (Geraerts *et al*., 2022; Huyse *et al*., 2015; Jorissen *et al*., 2022; Kmentová *et al*., 2019; Ondračková *et al*., 2012; Reyda *et al*., 2020). However, these studies focused on species-level invasions from the native to the non-native range, rather than intraspecific invasions of different lineages within the native range. Our study, on the other hand, demonstrates the use of parasites as tags to trace intraspecific cryptic invasions. Multiple instances of host lineage switches revealed by the phylogeographic analyses of *P. xenopodis* act as an indicator for historic contact between distinct *X. laevis* lineages. This provides us with the means of recognising such contact even in cases where the locally non-native lineage of *X. laevis* may have gone extinct after the introduction event or where they are not detected at a certain sampling effort.

In fact, thus far, genetic evidence for the admixture of the lineages of *X. laevis* with native ranges on opposite sides of the Great Escarpment has not been found, despite historical records pointing towards large scale exports from the southwest to the rest of southern Africa (van Sittert & Measey, 2016; Weldon *et al*., 2007). In our study, the host phylogeographic analyses suggest the presence of contact zones between the northeastern (NE) and southwestern lineages (SW_inland_ & SW_coast_) at two localities to the northeast of the Great Escarpment where these lineages co-occur.

It is this sympatry of multiple host clades that makes host lineage switching by parasite lineages a possibility. In turn, the observation of host switches from one lineage to another provides more evidence for the translocation of *X. laevis* across southern Africa. Indeed, our study confirmed the presence of southwestern parasite lineages (**sw1**, **sw2** & **sw3**) at seven localities to the northeast of the Great Escarpment, where one would expect the northeastern parasite lineage (**ne**) to occur. In these instances, southwestern parasite lineages that are translocated along with their respective host lineages, thus locally non-native at these new localities, spill over to the locally native northwestern host lineage when these divergent, previously allopatric host lineages come into contact (Galbreath & Hoberg, 2015).

Differential survival between locally native and non-native *X. laevis* lineages in southern Africa can partially explain the fewer observed instances of establishment by non-native *X. laevis* lineages when compared to its co-introduced polystomatid. In translocation experiments of the northern and southern lineages in southern Africa, *X. laevis* tadpoles and metamorphs exhibited remarkable phenotypic plasticity when translocated between rainfall regimes at the cost of longterm survival (Kruger, 2020). Under these circumstances, the locally native *X. laevis* lineage would outcompete the non-native lineage, preventing successful local invasion. Thus, the employment of host phylogeography alone to trace long range translocation of *X. laevis* in southern Africa would mask a number of introduction events, as our findings clearly demonstrate.

Upon arrival in the new area, the co-introduced *P. xenopodis* lineage may rapidly spread in a population of locally native and therefore naive *X. laevis*. It has been shown that primary infections by *P. xenopodis* in previously uninfected *X. laevis* elicit strong long term immunological responses against secondary infections by the same parasite species (Jackson & Tinsley, 2001). Correspondingly, a cross-infectivity experiment between northeastern and southwestern lineages of *X. laevis* and *P. xenopodis* further suggested that lineages of *X. laevis* in their native ranges might be more susceptible to infection by the locally non-native lineage of *P. xenopodis* than to their corresponding locally native parasite lineage (Jackson & Tinsley, 2005). In turn, this effect will boost the nascent spread of non-native *P. xenopodis* lineages in the locally native lineage of *X. laevis*.

There are two possible explanations for the observed contact between divergent lineages of this host-parasite system in its native range: natural geographic expansion over the evolutionary history of the host-parasite system in response to past environmental fluctuations, or more recent human-mediated translocation of the host along with its corresponding parasite lineage. At first glance, it is noticeable that the non-native *P. xenopodis* specimens collected at northeastern localities present with haplotypes that were generally highly differentiated from their counterparts from the southwest, despite belonging to the same **sw** lineages. In contrast, the haplotypes of the co-introduced French and Portuguese polystomatids are identical or nearly identical to the source South African haplotypes, indicating low divergence after the introduction event. The apparently deep divergence between native southwestern polystomatids and domestically translocated southwestern polystomatids suggests a scenario where parasites that underwent natural geographic expansion in deep time persist to the present day, yet with plenty of time to accrue genetic polymorphisms that differentiate them from their close relatives in the southwest. However, at our level of sampling effort, such comparisons of haplotype diversity are premature. It is clear from the long branch lengths between native southwestern polystomatids in the haplotype genealogy that there is considerable differentiation among the southwestern populations of *P. xenopodis*. It is reasonable to assume that greater sampling effort will yield more southwestern haplotypes that could be very similar to the translocated southwestern haplotypes now found at northeastern localities. Furthermore, it is worth noting that the number of domestic exports from the southwestern region of South Africa greatly dwarfs the number of the international exports (van Sittert & Measey, 2016), which translates to more introduction events of southwestern *X. laevis* to areas across South Africa, thus greater haplotype diversity among the domestically introduced frogs, than to France.

However, there is another reason affecting the plausibility of the scenario of earlier natural expansion of this host-parasite system. Of course, hosts and parasites can experience shared histories of geographic expansion and contact between isolated populations in response to historical environmental shifts, such as climate change (Galbreath & Hoberg, 2015; Hahn *et al*., 2015; Huyse *et al*., 2017; Kmentová *et al*., 2018). Likewise, post expansion, the non-native host lineages may be rapidly lost from the population due to genetic drift or differential survival, whilst the non-native parasite lineages may persist and become widely distributed across the geographic range of its new host lineage, as has been demonstrated before on the species level for tapeworm parasites in Beringia (Galbreath *et al*., 2020). Regardless, the formation of the Great Escarpment around 130 million years ago (Burke & Gunnel, 2008) predates the evolution of *X. laevis* over the last 15 million years (Furman *et al*., 2015). The Great Escarpment is a dominant barrier to dispersal that has played a defining role in shaping the distribution and evolution of southern African fauna (e.g. Barlow *et al*., 2013; Makokha *et al*., 2007; Mynhardt *et al*., 2015; Nielsen *et al*., 2018; Predel *et al*., 2012), not least among which *X. laevis* (Furman *et al*., 2015). When considering the evidence from the phylogeographic structuring of insects, frogs, lizards, snakes and small mammals in southern Africa, where the Great Escarpment is a definite barrier to natural dispersal (Barlow *et al*., 2013; Furman *et al*., 2015; Makokha *et al*., 2007; Mynhardt *et al*., 2015; Nielsen *et al*., 2018; Predel *et al*., 2012), it seems highly unlikely that climatic fluctuations could have given rise to natural expansion of *X. laevis* beyond the plateau edge of the escarpment that may shape the distribution of the southwestern lineages of *P. xenopodis* to the current day.

Rather, we consider recent human-mediated translocation a more plausible explanation. In fact, we know that the bulk of exports of *X. laevis* since the 1930s from the southwest of Southern Africa, which amounts to over 400 000 animals, was sent to domestic destinations, especially to urban centers inland where *X. laevis* was used in research and teaching (van Sittert & Measey, 2016; Weldon *et al*., 2007). Subsequent escapees from these centers of higher learning would lead to the presence of locally non-native *X. laevis* and *P. xenopodis* near these urban areas. Moreover, these escapees can spread even further through the ability of *X. laevis* to move overland, a mode of dispersal which is complemented by expansion via networks of artifical waterways (de Villiers & Measey, 2017; Measey, 2016). The rural locations of the parasite spillover events revealed by our analyses of polystomatid phylogeography suggest that smaller scale human-mediated dispersal events also have a role to play in the dispersal of *X. laevis*. The role of recreational fishing across South Africa in the dispersal of *X. laevis*, and by extension, *P. xenopodis*, has been recognised before (Weldon *et al*., 2007). Previously documented interviews with local anglers and suppliers exposed the use of *X. laevis*, frequently bought from pet shops and local breeders, as live bait for predominantly catfish angling (Weldon *et al*., 2007). Our own informal discussions with local anglers at the localities where we collected *X. laevis* affirm the widespread use of this frog, sometimes obtained from pet shops, as live bait for the capture of both black bass (*Micropterus* spp.) and catfish (*Clarias gariepinus*).

In terms of sheer numbers, the impact of these translocation events on the domestic dispersal of *X. laevis* has been deemed negligible when compared to exports from Jonkershoek, the most prolific exporter, and other official suppliers to the domestic market (van Sittert & Measey, 2016; Weldon *et al*., 2007). Nonetheless, previous studies on other aquatic species have pointed out that the use of live bait in angling is an important pathway for introductions, even if the market might be small in economic terms (Kalous *et al*., 2013; Kilian *et al*., 2012). Therefore, the impact of anglers on the spread of locally non-native *X. laevis* lineages to more remote areas, as opposed to the bigger centra that received the documented official consignments, should not be underestimated. Our informal interviews identified a myriad of ways for *X. laevis* to be released in new environments by anglers, from escapees *en route*, to the deliberate release of surplus bait, to the intentional stocking of water bodies for future use. Therefore, the combination of overland movement, dispersal along natural and artificial waterways and translocation by the angling community could form a widespread undocumented dispersal network of *X. laevis* in South Africa.

### 4.3 Parasite phylogeography as an underutilised tool in invasion biology

As shown above, the present study acts as a proof of concept for the potential of the comparative phylogeographic approach to the clarification of patterns in intraspecific host divergence and dispersal. In turn, this independent line of evidence can provide additional insight on invasion pathways and inform management strategies. Here, we highlight some of the potential implications to demonstrate the broader applicability of this integrative approach.

Our approach was able to provide a new perspective on the origin of the relatively high haplotype diversity of the invasive population of *X. laevis* in France, recovered in our analysis and others, when compared to other invasive populations in Portugal, Sicily and Chile (de Busschere *et al*., 2016; Lillo *et al*., 2013; Lobos *et al*., 2014). The idea of multiple introduction events from various source populations in South Africa has been offered before as the most parsimonious explanation for this genetic diversity in the invasive population of *X. laevis* in western France (de Busschere *et al*., 2016). Since we found evidence for host switches by *P. xenopodis* in the native range, an event that necessarily involves the sympatry of divergent host lineages, it is a certainty that multiple host lineages co-occur at certain localities in the native range. In this, our results build upon the evidence provided by historical export records (van Sittert & Measey, 2016; Weldon *et al*., 2007), dispersal patterns (de Villiers & Measey, 2017; Measey, 2016) and anecdotal information (Weldon *et al*., 2007) to show that the mixing of lineages most likely already happened in the source populations in South Africa prior to the co-introduction of *X. laevis* and *P. xenopodis* to western France (van Sittert & Measey, 2016; Weldon *et al*., 2007).

The novel insights provided by the addition of the parasite data set has implications for both the conservation of *X. laevis* and its host-specific parasites. The widespread anthropogenic translocation of this host-parasite system disrupts interactions between *X. laevis* lineages and the *P. xenopodis* lineages that tracked their phylogenetic divergence, ancient interactions that deserve to be targets of conservation (Gómez & Nichols, 2013). Subsequent host switches between closely related lineages set the stage for new diversity to arise and for novel ecological interactions to be established (Galbreath *et al*., 2020; Sures *et al*., 2017). In the case of host-specific parasites, translocation of lineages within the native range of a species may actually be of more concern than translocation to far-flung non-native ecosystems, since these ecosystems will probably not harbour close relatives of the invasive host to facilitate host switches (Blackburn & Ewen, 2017). Host-parasite interactions are also something to keep in mind when conducting conservation re-introduction programmes, where captive-bred or locally non-native individuals of a species are used to re-stock locally extinct populations of endangered species (Britt *et al*., 2004). Our results highlight that the evaluation of the genetic and parasitological suitability of a source population for translocation should also encompass the potential history of local introductions that may have influenced the intraspecific parasitism of the host population (Jørgensen, 2015; Northover *et al*., 2018).

This approach further holds great promise in the management of invasive species. Prior to invasion, translocations in the native range may have already altered the source populations of invasive species, as is the case of *X. laevis*. We need to take altered phylogeographic structure into account to adequately monitor the course of an invasion or the novel interactions that may arise. Parasites as passengers or invaders themselves easily slip under the radar in invasion biology (Blackburn & Ewen, 2017; Solarz & Najberek, 2017). Yet, the role of parasites on the move in emerging infectious diseases and habitat transformation via indirect density-mediated effects cannot be overemphasised (Amundsen *et al*., 2013; Bobadilla-Suarez *et al*., 2017; Daszak, 2000; Dunn *et al*., 2012; Goedknegt *et al*., 2016; Lymbery *et al*., 2014; Roy & Lawson Handley, 2012; Sures *et al*., 2017). Furthermore, the identification of the local angling community as a player in the translocation of *X. laevis* in the native range allows policy makers to broaden their intervention efforts in the management of *X. laevis* as a domestic exotic.

### 4.4 Recommendations to make parasite data sets work for invasion biologists

All in all, our findings agree with the necessity of a more holistic approach to data collection in invasion biology (Clavero *et al*., 2016), especially with regard to parasites. The idea of the ”holistic specimen” has recently gained some traction (Cook *et al*., 2017; Galbreath *et al*., 2019). This approach to the collection and study of organisms advocates for the integrated specimen-based studies of animals and their associated parasites and pathogens. The optimisation of sampling effort is crucial to improve the allocation of resources, both in terms of funding and specimen use, especially when working with endangered species or specimens that require considerable effort to obtain, or in view of other ethical concerns regarding invasive sampling.

The inclusion of parasite data during the data collection phase also ensures that the host and parasite data sets are comprised of directly comparable individuals which can improve the feasibility and value of comparative phylogeographic analyses. For example, the non-overlap between host and parasite data sets was cited as a limitation in the co-phylogeographic analyses conducted by both Barbosa *et al*. (2012) and Huyse *et al*. (2017). Overlap between data sets can be established through the sampling of host-parasite pairs from the same localities and time frames, whilst sampling of the corresponding parasite of each host specimen will not contribute to data set overlap, especially in the case of parasites with a free-living stage. Even in cases where parasites are not the targets of the current project, the deposition of voucher material of both host and parasite from the sampling event into curated institutional collections, in other words overlapping data sets, can be vital for later studies (Galbreath *et al*., 2019; Thompson *et al*., 2021). Later on, DNA sequences could be linked to material deposited in these museum collections.

Similarly, another aspect to keep in mind is the choice of markers for our co-phylogeographic analyses. More often than not, investigators rely on markers for which primers are already available and entire analyses may even be based on a single marker (de Vienne *et al*., 2013; Gutiérrez-García & Vásquez-Domínguez, 2011). Yet, successful comparative phylogeographic analyses rely on a combination of multiple markers with a sufficient degree of genetic divergence evident among populations (de Vienne *et al*., 2013; Gutiérrez-García & Vásquez-Domínguez, 2011), which is often higher in the host than the parasite (Nieberding & Olivieri, 2007). Thus, the independent yet comparable DNA signature provided by the parasite data set may provide higher resolution than the host data set, where spatial patterns may be blurred by the recent mixing of previously isolated genotypes in invasive populations (Rius & Turon, 2020). As a next step to improve confidence in phylogeographic inferences, we also propose the addition of morphological data from the parasite specimens, as demonstrated by several recent studies on monogenean parasites (Huyse *et al*., 2015; Kmentová *et al*., 2019; Ondračková *et al*., 2012; Schoeman *et al*., 2022).

Ultimately, this study serves as a proof of concept that the use of parasite data sets will add value to investigations in invasion biology, especially in the case of intraspecific cryptic invasions. We show that the utility of parasite data sets can go beyond the provision of supporting evidence and provide greater resolution that may lreveal patterns that went undetected in the host data set. Our study echoes the sentiments of Hoberg *et al*. (2015), who argued for an integrated approach to parasitology as a discipline that holds the keys to understanding and managing global change in the Anthropocene.

## DATA AVAILABILITY STATEMENT

All DNA sequences used for co-phylogeographic analyses will be made available on the public database GenBank, along with relevant metadata and accession numbers will be added upon article acceptance. Detailed locality data are contained in the supplementary information.

## ACKNOWLEDGMENTS

ALS received funding from the DST-NRF Centre of Excellence for Invasion Biology (South Africa). The project utilised the infrastructure of the Unit for Environmental Sciences and Management, North-West University (South Africa). This research was supported by EMBRC Belgium – FWO project GOH3817N. NK and MPMV are financed by the Special Research Fund of Hasselt University (BOF21PD01 and BOF20TT06, respectively). The authors would like to express their sincere thanks to a number of persons who assisted in the collection of the frogs. Guénolé le Peutrec, Jean Secondi, Natasha Kruger and Rodolphe Olivier helped with the collection in France. Rui Rebelo provided access to frogs from Portugal. In South Africa, several farm owners graciously gave permission for collection to take place on their properties and provided lodging for the research team: Fanus and Olga Kritzinger, William and Christa van Zyl, Dave Schlebusch, Fanus and Carin Oberholzer, Danie and Annalise Marais, Johan Hamann, Tobie Bielt, Gert Bench, Stoffel Labuschagne, Jannie and Susan van Rensburg, Jan Meintjies, Marthinus Hartman, Douw and Louise de Jager and Ernest de Villiers. Lastly, Mathys Schoeman, Clarke Scholtz, Andrea Darvall, Willie Landman, Ferdi de Lange, John Measey and Roxanne Viviers assisted with the collection of frogs at the remainder of the localities in South Africa. The authors are also indebted to numerous local recreational fishermen who were willing to share details about their use of *Xenopus laevis* across South Africa. The utilisation of the frogs and the research protocols were approved by the Animal Care, Health and Safety in Research Ethics (AnimCare) Committee of the Faculty of Health Sciences of the North-West University (ethics number: NWU-0380-16-A5-01). Animals from the South African populations were sampled under the permits 0056-AAA007-00224 (CapeNature) and FAUNA 1343-2017 (Northern Cape) provided by the Department of Economic Development, Environmental Affairs and Tourism.

## AUTHORS’ CONTRIBUTIONS

ALS conceived the initial idea, collected and analysed the data and led the writing of the manuscript. All authors refined the initial concept, designed the methodology, contributed to interpretation of data, contributed critically to the drafts and gave final approval for publication.

