## Supplemental Table S1 for "A monogenean parasite reveals the widespread translocation of the African Clawed Frog in its native range"

### Appendix S1: Supplementary information on the collection localities of *Xenopus laevis* for ”A monogenean parasite reveals the widespread translocation of the African Clawed Frog in its native range”

**Anneke L Schoeman, Louis H du Preez, Nikol Kmentová and Maarten PM Vanhove**

**Table S1.1.** Detailed information on collection localities from continents where *Xenopus laevis* and their corresponding *Protopolystoma xenopodis* specimens were sampled for this study. Included are 59 specimens for co-phylogeographic analysis from 42 water bodies from western France, Portugal and southern Africa. Localities in southern Africa are named according to the nearest town and province. Water bodies in southern Africa, France and Portugal were further classified according to the phylogeographic lineage of the collected *X. laevis* (SA1–7), following de Busschere *et al.* (2016) (see Appendix S2 for further details). In addition, for each surveyed water body, collection month (Date), latitude and longitude in decimal degrees (Coordinates) and number of host-parasite pairs (Sample size) are given.

| Locality | Lineage | Date | Coordinates | Sample size |
|---|---|---|---|---|
| **Southern Africa** | | | | |
| Adelaide, Eastern Cape | SA6 | November 2017 | 32.6879°S; 26.2951°E | 2 |
| Barrydale, Western Cape | SA3 | November 2017 | 33.9264°S; 20.5917°E | 1 |
| | | | 33.9283°S; 20.5915°E | 2 |
| Cape Town, Western Cape | SA1 | June 2017 | 33.8355°S; 18.5528°E | 1 |
| Debshan ranch, Zimbabwe | SA5 | November 2017 | 19.6825°S; 29.1547°E | 1 |
| Dullstroom, Mpumalanga Province | SA5 | April 2017 | 25.3981°S; 30.0380°E | 1 |
| Durbanville, Western Cape | SA1 | July 2017 | 33.8392°S; 18.6003°E | 2 |
| George, Western Cape | SA2 | February 2018 | 33.9455°S; 22.4637°E | 2 |
| Hermanus, Western Cape | SA1 | June 2017 | 34.3702°S; 19.2571°E | 1 |
| Hopetown, Northern Cape | SA5 | December 2017 | 33.3528°S; 20.4255°E | 1 |
| Johannesburg, Gauteng Province | SA5 | February 2018 | 26.1251°S; 27.9222°E | 1 |
| Jonkershoek, Western Cape | SA1 | February 2020 | 33.9584°S; 18.9169°E | 1 |
| Ladismith, Western Cape | SA3 | November 2017 | 33.4637°S; 21.0970°E | 3 |
| Laingsburg, Western Cape | SA3 | November 2017 | 33.3528°S; 20.4255°E | 2 |
| Louis Trichardt, Limpopo Province | SA5 | February 2019 | 23.0728°S; 30.0711°E | 1 |
| Matjiesfontein, Western Cape | SA3 | February 2020 | 33.2294°S; 20.5816°E | 3 |
| Modimolle, Limpopo Province | SA5 | April 2017 | 24.6384°S; 28.4369°E | 1 |
| Potchefstroom, North-West Province | SA5 | March 2017 | 26.7555°S; 27.0506°E | 1 |
| Riversdale, Western Cape | SA3 | November 2017 | 33.7033°S; 21.1762°E | 1 |
| Three Sisters, Western Cape | SA4 | November 2017 | 31.8950°S; 23.0957°E | 3 |
| | | | 31.8526°S; 23.2645°E | 5 |
| Tzaneen, Limpopo Province | SA5 | May 2017 | 23.7988°S; 30.1951°E | 1 |
| Ugie, Eastern Cape | SA6 | December 2017 | 31.3049°S; 28.2646°E | 1 |

**Table S1.1.** Detailed information on collection localities where *Xenopus laevis* and their corresponding *Protopolystoma xenopodis* specimens were sampled for this study (*continued*).

| Locality | Lineage | Date | Coordinates | Sample size |
|---|---|---|---|---|
| Vanrhynsdorp, Western Cape | SA7 | November 2017 | 31.7361°S; 18.8255°E | 3 |
| Verkykerskop, Free State | SA5 | January 2018 | 27.7833°S; 29.3363°E | 1 |
| Wakefield, KwaZulu Natal | SA6 | February 2018 | 29.4797°S; 29.8877°E | 1 |
| | | April 2019 | 29.4825°S; 29.8889°E | 1 |
| Wellington, Western Cape | SA1 | November 2017 | 33.5714°S; 18.8401°E | 3 |
| White River, Mpumalanga Province | SA5 | April 2017 | 25.3391°S; 31.0226°E | 1 |
| | | | 25.3320°S; 31.0433°E | 1 |
| **France** | | | | |
| Sites 1, 3, 5 | FR | July 2017 | 47.3440°N; 0.7636°W | 1 |
| | | | 46.9401°N; 0.3686°W | 1 |
| | | | 47.0328°N; 0.3474°W | 1 |
| Ambillou-Château | FR | June 2019 | 47.2386°N; 0.3445°W | 1 |
| Bouillé-Saint-Paul | FR | June 2019 | 47.0273°N; 0.3443°W | 1 |
| Brézé | FR | June 2019 | 47.1726°N; 0.0729°W | 1 |
| Cersay | FR | May 2019 | 47.0531°N; 0.3648°W | 1 |
| | | July 2019 | 47.0531°N; 0.3648°W | 1 |
| Chalonnes-sur-Loire | FR | June 2019 | 47.3440°N; 0.7636°W | 1 |
| Le Coudray-Macouard | FR | June 2019 | 47.1872°N; 0.1151°W | 1 |
| Massais | FR | June 2019 | 47.0106°N; 0.3582°W | 1 |
| Saint-Martin-de Sanzay | FR | May 2019 | 47.0729°N; 0.1898°W | 1 |
| **Portugal** | | | | |
| Lisbon | PT | February 2019 | 38.7136°N; 9.2897°W | 1 |
