## Supplemental Table S2 for "A monogenean parasite reveals the widespread translocation of the African Clawed Frog in its native range"

### Appendix S2: Supplementary information on the classification of collection localities for ”A monogenean parasite reveals the widespread translocation of the African Clawed Frog in its native range”

**Anneke L Schoeman, Louis H du Preez, Nikol Kmentová and Maarten PM Vanhove**

There is remarkable genetic and morphological divergence among different populations of *X. laevis* in southern Africa (du Preez *et al.*, 2009; Furman *et al.*, 2015; Grohovaz *et al.*, 1996; Measey & Channing, 2003). In the present study, the collection localities are classified according to their proximity to previously sampled localities where the phylogeographic lineages of the *X. laevis* specimens have been determined, retrospectively named by de Busschere *et al.* (2016) (Figure S2.1).

Accordingly, *X. laevis* collected along the coast to the southwest of the Cape Fold Mountains were assigned to SA1, or SA2 if these coastal localities were more to the east. The SA3 *X. laevis* were captured among the Cape Fold Mountains near the Laingsburg contact zone between the southwestern coast and the Great Escarpment to the northeast, with SA4 *X. laevis* captured further northward in the lowland Karoo semi-desert in the summer rainfall region near the plateau edge of the Great Escarpment. The *X. laevis* captured to the north of the Cape Fold Mountains near the western coast in the winter rainfall region were designated SA7. Finally, *X. laevis* from the summer rainfall region to the northeast were classified as SA5 if they were captured on the inland plateau of the Great Escarpment or as SA6 if they were captured at the edge of the Great Escarpment along the eastern seaboard.

The South African localities were also divided into southwestern and northeastern localities according to their position relative to the Great Escarpment for the calculation of genetic diversity and interpretation of the haplotype networks. Accordingly, the SA1, SA2, SA3 and SA7 localities are regarded as southwestern and the SA5 and SA6 localities are regarded as northeastern. The SA4 locality is a known contact zone between the two, but was regarded as a southwestern locality for the calculation of genetic diversity of the predefined groups.

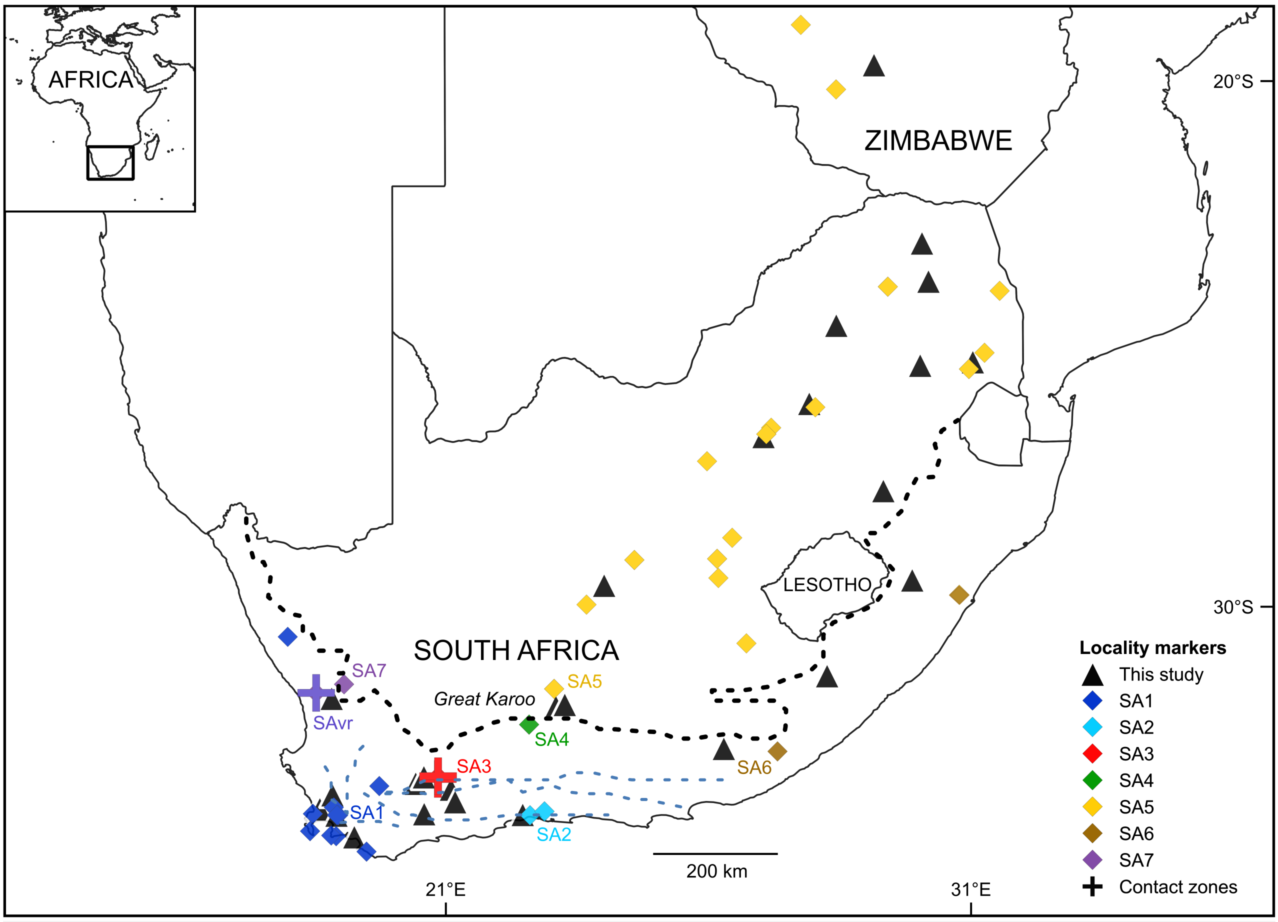


**Figure S2.1.** The phylogeography of *Xenopus laevis sensu stricto* across southern Africa. The localities (◆) where *X. laevis* were collected by de Busschere *et al.* (2016); du Preez *et al.* (2009); Furman *et al.* (2015); Grohovaz *et al.* (1996); Measey & Channing (2003) are coloured according to the frogs' phylogeographic lineage proposed by the respective authors, retrospectively named SA1, SA2, SA3, SA4, SA5, SA6 and SA7 by de Busschere *et al.* (2016). The localities sampled for the present study (▲) are also included, along with the previously identified contact zones (✚), namely SA3 and SAvr. The Great Escarpment (black dotted line) and the Cape Fold Mountains and associated mountain ranges (blue dotted lines) were derived from the Natural Earth database. The map was constructed in QGIS *version* 3.10.2-A Coruña (QGIS Development Team, 2018) with the Mercator projection.
